# EM3DFold: accurate de novo protein and nucleic acid model building for cryo-EM maps using language model-powered deep learning

**DOI:** 10.64898/2026.09.16.752067

**Authors:** Tao Li, Hong Cao, Sheng-You Huang

## Abstract

Cryo-electron microscopy (cryo-EM) has become one of the most powerful techniques for macromolecular structure determination. However, accurate model building from cryo-EM maps remains challenging, particularly for nucleic acids. Here, we present EM3DFold, a unified *de novo* model-building framework for accurate structure determination of proteins, nucleic acids, and protein-nucleic acid complexes from cryo-EM maps using a density-aware, large language model-powered three-track attention (TTA) network. The TTA network effectively integrates sequence, density, and structural information to enable accurate all-atom model building. EM3DFold was extensively evaluated on independent benchmarks of 298 experimental cryo-EM maps at *<* 4.0 Å resolutions, and achieves an unprecedentedly high median accuracy of 75% completeness (88% coverage and 94% sequence accuracy) for 178 nucleic acid maps, 90% completeness (95% coverage and 96% sequence accuracy) for 124 protein-nucleic acid complexes, and 95% completeness (97% coverage and 98% sequence accuracy) for 120 protein-only targets, substantially outperforming state-of-the-art methods including ModelAngelo, EM2NA, CryoREAD, and EMProt. In addition, EM3DFold also produces models with superior model-to-map fit and stereochemical quality, providing a robust and reliable solution for automated cryo-EM model building. The EM3DFold package is freely available at https://github.com/huang-laboratory/EM3DFold.

## 1 Introduction

Proteins and nucleic acids are fundamental to many biological processes, functioning either independently or through intricate interactions with other macromolecules. Consequently, accurate determination of their three-dimensional atomic structures is indispensable for elucidating the molecular mechanisms underlying life processes. Over the past decade, cryo-electron microscopy (cryo-EM) has transformed the field of structural biology, emerging as a powerful technique capable of visualizing biological macromolecules in near-native states at near-atomic resolution^1–3^. Advances in hardware and software have increasingly accelerated this revolution^4–9^, enabling routine determination of structures that were previously intractable. As a result, the Electron Microscopy Data Bank (EMDB)^10^ has experienced an exponential increase in cryo-EM map depositions. Furthermore, interpreting these high-resolution maps to atomic models and depositing them in the Protein Data Bank (PDB)^11^ greatly enrich the structural knowledge base for a wide range of macromolecular complexes.

Building accurate atomic models from cryo-EM density maps requires both expertise and specialized computational tools. Recent years have seen the fast development of automated protein structure determination from cryo-EM maps—either via *de novo* modeling or structure fitting^12–26^. However, methods for model building of nucleic acids from cryo-EM maps are still limited, owing to the challenges in distinguishing similar bases and low resolutions of their cryo-EM maps. As a representative of earlier *de novo* methods, phenix.map to model^27^, marks a major step for automated map-to-model studies. It integrates multiple processing pipelines of the Phenix^7^ framework and shows acceptable precision for backbone modeling of proteins and nucleic acids. DRRAFTER^28^ and its successor auto-DRRAFTER^29^ can model nucleic acids with known secondary structures, but depend on extensive conformational sampling and demand substantial computational time and resources.

Recently, artificial intelligence (AI), like deep learning (DL), has been used to address the modeling challenges in the determination of nucleic acid structures from cryo-EM maps. ModelAngelo^19^, as a leading approach, achieves a high accuracy in modeling proteins, but suffers from low sequence accuracy in nucleic acid model building. Other DL-based methods like EMRNA^30^, EM2NA^31^, and CryoREAD^32^ have also been proposed. For these methods, deep neural networks are first used to predict positions of main-chain atoms and nucleotide types, followed by post-processing such as heuristic algorithm-based fragment tracing and sequence alignment for final model building. These DL-based methods achieve reasonable atomic position predictions with high computational efficiency, but they all have various limitations in structure quality, model-to-map fit, and/or sequence assignment.

Despite the present progresses, model building of cryo-EM maps still faces challenges such as poor backbone RMSD, atomic clashes, and low sequence accuracy at near-atomic resolutions, especially for protein-nucleic acid complexes. Addressing the challenges, we present EM3DFold, a universal framework for accurate *de novo* model building of proteins, nucleic acids, and protein-nucleic acid complexes from cryo-EM maps. EM3DFold features several key components: a main-chain atom predictor based on the Swin-Conv-UNet (SCUNet)^33^ network, an all-atom and residue type predictor based on a density-aware Three-Track Attention (TTA) neural network, and post-processing algorithms for all-atom modeling. The TTA network design enables the efficient integration of sequence, density, and structure. The core module of our pipeline for the all-atom predictor enables accurate and efficient determination of full atomic coordinates directly from local density signals.

EM3DFold was extensively evaluated on independent benchmarks of 298 cryo-EM maps, including 54 nucleic acid-only targets, 124 protein-nucleic acid complexes, and 120 protein-only structures with resolutions better than 4 Å, and was compared with state-of-the-art automated model-building methods, including EM2NA, CryoREAD, ModelAngelo, and EMProt. It is shown that EM3DFold consistently outperforms existing approaches. In addition, the models generated by EM3DFold also exhibit superior model-to-map agreement and structure quality, demonstrating its high ability to produce complete and reliable atomic models.

## 2 Results

### 2.1 The workflow of EM3DFold

Figure 1 shows the workflow of EM3DFold for both protein and/or nucleic acid model building from a cryo-EM map. The inputs of EM3DFold are the target density map and sequences (optional). First, the density map is fed into the main-chain C*α*/C4’ atom predictor to get a C*α*/C4’ atom probability map. We then convert the atom probability map to C*α*/C4’ atom coordinates by detecting local maxima using a mean-shift algorithm. The C*α*/C4’ atoms are input to the all-atom refiner for predicting the refined C*α*/C4’ positions, the backbone frames, the torsion angles, and the residue types. With the C*α*/C4’ atoms, backbone frames, and torsion angles, we can derive the backbone atom positions by transforming ideal amino acid/nucleotide atom groups^34, 35^. Since the resulted main-chain atoms are unordered, to make nucleotides form chains, we then thread individual residues to obtain backbone traces by iteratively searching the next nearest residue.

**Figure 1:**
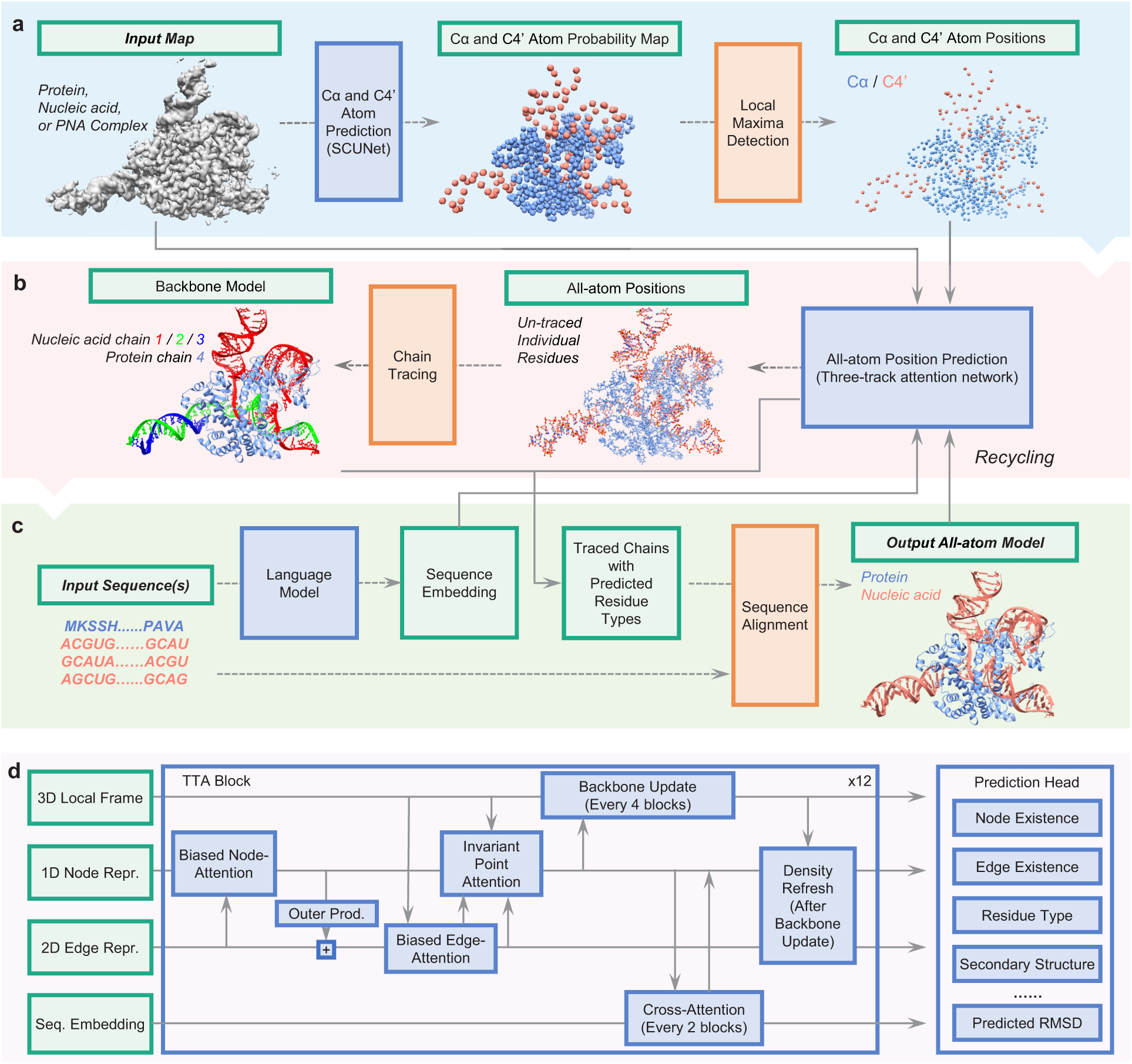
Overview of the EM3DFold framework. The workflow of EM3DFold can be divided into three stages. **a**, The C*α* and C4’ atom prediction stage. In this stage, the input density map is processed by a trained SCUNet to predict C*α* and C4’ probability map, from which C*α* and C4’ positions are extracted using mean-shift clustering. **b**, The backbone modeling stage. In this stage, the C*α*/C4’ positions are passed to a Three-Track Attention (TTA) network to predict backbone frames, torsion angles and residue types, enabling the placement of backbone atoms. Residues are then threaded to backbone fragments using O3’–P and C–N bond length restraints. **c**, The sequence assignment stage. In this stage, the predicted residue type logits of the backbone structures are used to construct HMMs and then are searched against the target sequences to assign residue types. The pipeline also applies extra recycling steps for model refinement before the final model is output. **d**, Architecture of the TTA module, where 1D, 2D, and 3D features and target sequence embeddings interact through specialized attention mechanisms. EM3DFold adopts multiple prediction tasks during training to encourage model to learn local structure.

Next step is the determination of residue (amino acid or nucleotide) types, the major challenge for existing modeling methods, especially on nucleic acids. EM3DFold adopts two residue type predictors. One is a SCUNet-based^33^ residue type predictor (only for nucleic acids) that leverages only density map information, which is powerful for density maps at high resolution. The other is a Three-Track Attention (TTA) network predictor that incorporates both density and sequence information. In addition to predicting atom positions, the TTA network also outputs the residue type for each node. This design enables the network to jointly learn structural relationships and predict plausible residue types. We also incorporate sequence embeddings from large language models, including ESM^36^ for proteins and RiNALMo^37^ for nucleic acids, into the TTA networks.

With the predicted residue types, we construct two HMM files for each backbone fragment: one is from the TTA-based predictor and the other is from the SCUNet-based predictor. Each HMM is searched against the target sequences to find the best matched parts using HMMER^38, 39^. The HMM file with the higher match score is kept for the fragment. The corresponding sequence alignment is used to assign the amino acid/nucleotide types for the backbone fragment. With the assigned residue types, the positions for the side-chain atoms are finally determined using the predicted torsion angle. Similar to that in AlphaFold^40, 41^, the structure is fed to the networks again for another round of modeling. A final structure is output after four recycling steps.

### 2.2 Performance on 178 nucleic acid maps

EM3DFold is first evaluated and compared with three state-of-the-art nucleic acid model building methods, including EM2NA, CryoREAD, and ModelAngelo, on the 178 nucleic acid maps from 54 nucleic acids and 124 protein-nucleic acid complexes. Four metrics are used to measure model accuracy: backbone RMSD, residue coverage, sequence accuracy, and completeness. Among these metrics, backbone RMSD and residue coverage reflect the accuracy of backbone modeling without considering residue types, while sequence accuracy assesses the proportion of residues that are correctly determined in both backbone geometry and residue type. Completeness is the product of coverage and sequence accuracy, representing the overall accuracy of model building.

Figure 2a-d shows the comparison between EM3DFold and other methods in modeling nucleic acids. It can be seen from the figure that EM3DFold greatly outperforms the other three methods in all metrics. For the atomic accuracy of built models, EM3DFold obtains the lowest median backbone RMSD of 1.05 Å, compared with 1.70 Å for EM2NA, 4.15 Å for CryoREAD, and 1.22 Å for ModelAngelo (Figure 2a). In addition, EM3DFold and EM2NA achieve the best coverage of 88%, compared with 50% for CryoREAD and 84% for ModelAngelo (Figure 2b). In particular, EM3DFold achieves an excellent sequence accuracy of 94%, which is drastically higher than 55% for EM2NA, 56% for CryoREAD, and 43% for ModelAngelo (Figure 2c). Consequently, EM3DFold obtains the highest completeness of 75%, compared with 45% for EM2NA, 26% for CryoREAD, and 30% for ModelAngelo (Figure 2d). These results suggest that the improvement of EM3DFold over other methods comes from sequence assignment more than residue coverage of built models. The better performance of EM3DFold than existing methods is also observed in the refined models, where the refinement slightly improves the backbone RMSD but has little effect on coverage, sequence accuracy, and completeness.

**Figure 2:**
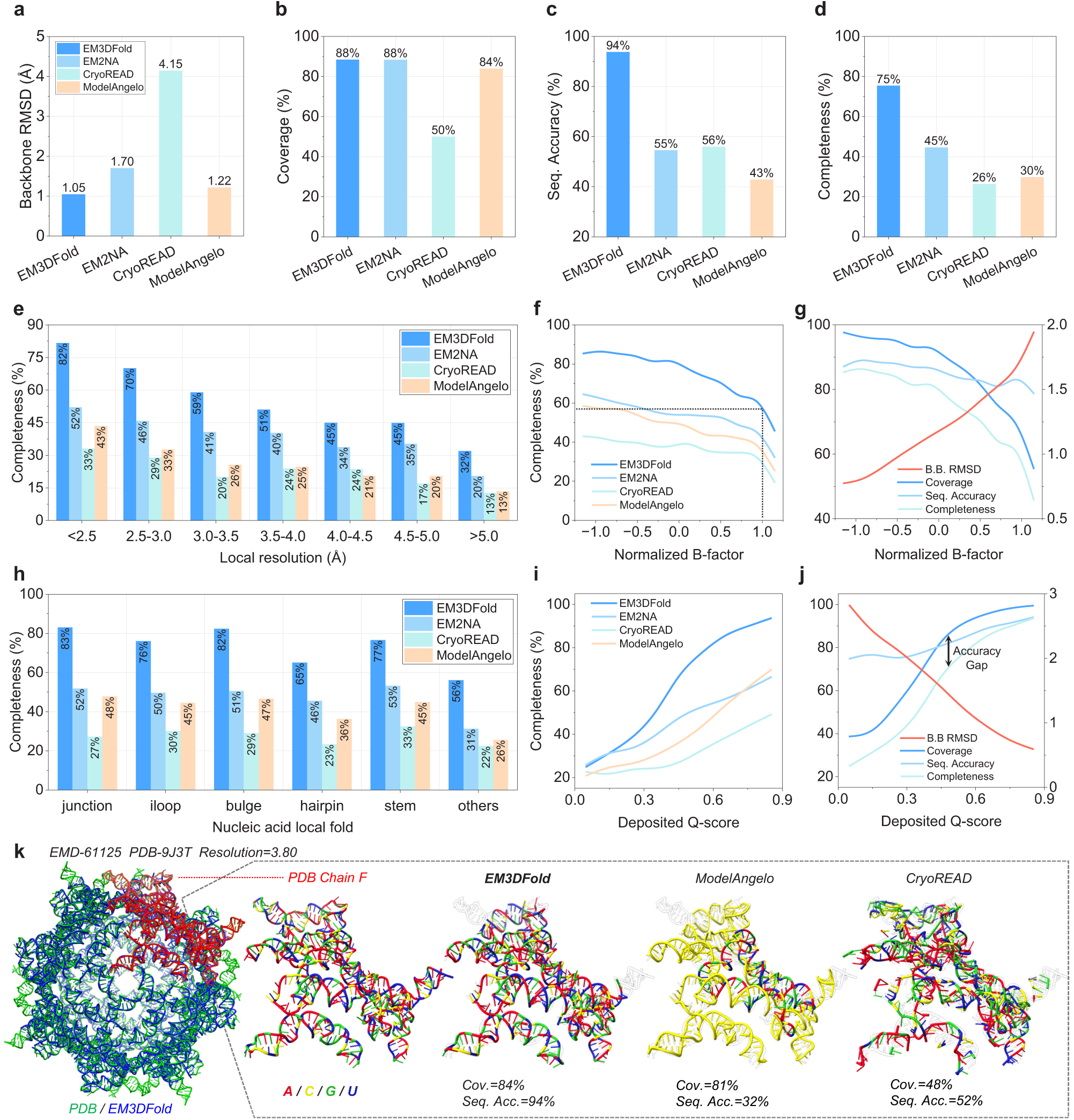
Performance of EM3DFold and other methods on *n* = 178 nucleic acid maps. **a-d**, Comparison of EM3DFold, EM2NA, CryoREAD, and ModelAngelo on nucleic acid modeling performance, including (**a**) backbone RMSD, (**b**) residue coverage, (**c**) sequence accuracy, and (**d**) completeness. Bars indicate medians. **e**, Completeness comparison for different local resolution ranges. **f**, Completeness as a function of normalized B-factor for EM3DFold and other methods. **g**, Backbone RMSD, residue coverage, sequence accuracy, and completeness of EM3DFold versus normalized B-factor. Left y-axis is for residue coverage, sequence accuracy, and completeness. Right y-axis refers to backbone RMSD. **h**, Completeness comparison for different nucleic acid local fold types, including junction, internal-loop (iloop), bulge, hairpin, stem, and other regions. **i**, Completeness as a function of deposited Q-score for different methods. **j**, Backbone RMSD, residue coverage, sequence accuracy, and completeness of EM3DFold versus deposited Q-score. Left y-axis is for residue coverage, sequence accuracy, and completeness. Right y-axis refers to backbone RMSD. **k**, Representative example (Enterococcus faecalis ROOL RNA octamer, PDB-9J3T, EMD-61125) of RNA modeling by EM3DFold, EM2NA, and CryoREAD. The PDB structures is shown in green, and the EM3DFold models in blue. The enlarged views show the subunit chain F modeled by different methods, with nucleotides colored according to residue types. On this target, EM3DFold achieves the highest sequence accuracy and completeness than the other methods.

To evaluate the robustness of automated modeling under heterogeneous density quality, we analyzed the model performance with respect to local resolution. Local resolution was estimated using ResMap^42^ for cryo-EM maps with available half-maps deposited in the EMDB. Figure 2e shows the completeness of the built models by different methods across different local resolution ranges. It is shown that EM3DFold consistently achieves higher completeness than the other approaches across all resolution ranges, with more advantages in lower resolution regions, e.g. *>* 3.5 Å. The performance improvement is also evident at higher resolutions, where EM3DFold yields 82% completeness for maps with local resolution better than 2.5 Å, compared with 52% for EM2NA, 33% for CryoREAD, and 43% for ModelAngelo. These results demonstrate the robustness of EM3DFold in building more complete models, particularly for structures with heterogeneous density.

To investigate the impact of structure flexibility (uncertainty), we analyzed the reported metrics with respect to residue B-factor (Z-score normalized) and Q-score^43^ calculated with the deposited PDB model. Figure 2f-g displays the model completeness across different normalized B-factors. It can be seen from Figure 2f that EM3DFold consistently achieves higher completeness than other methods across the entire B-factor range. Notably, as the B-factor increases, the completeness of all methods gradually decreases; however, EM3DFold maintains a substantial advantage over the other approaches. Even in regions with high normalized B-factors of 1.0, EM3DFold preserves approximately 57% completeness, compared with 45%, 35%, and 30% for EM2NA, CryoREAD, and ModelAngelo, respectively. In particular, with increasing B-factor, despite the sharply degradation in backbone RMSD, coverage and completeness, EM3DFold maintains high sequence accuracy of above *∼*80% (Fig. 2g).

Figure 2i-j shows the model completeness across different ranges of deposited Q-scores. Again, EM3DFold consistently achieves higher completeness across the entire Q-score range, demonstrating a substantially stronger modeling robustness again density quality. In particular, EM3DFold reaches over 90% completeness at high Q-scores, whereas EM2NA, CryoREAD, and ModelAngelo remain below approximately 70% (Figure 2i). This suggests that EM3DFold can more effectively exploit high-quality density information to generate complete nucleic acid models. In addition, similar to the trend versus normalized B-factors, sequence accuracy remains high across different Q-score ranges (Figure 2j). It is also observed that there is approximately a 15% gap between coverage and sequence accuracy across all Q-score ranges, indicating that sequence assignment remains a common bottleneck for automated model building (Figure 2j).

Figure 2h compares the completeness of generated models for different nucleic acid local fold types, including junctions, iloops, bulges, hairpins, stems, and other regions (annotated by x3dna^44^). It can be seen from the figure EM3DFold achieves the highest completeness across all structural categories, demonstrating its superior capability in modeling diverse nucleic acid architectures. The improvement is particularly evident for complex and flexible local folds. Even for stem regions, where density interpretation is generally easier, EM3DFold maintains a clear advantage with 77% completeness. These results indicate that EM3DFold provides more robust model building across different nucleic acid structural motifs, particularly for challenging non-canonical RNA folds.

Figure 2k presents an example of nucleic acid modeling on the *Enterococcus faecalis* ROOL RNA octamer (PDB: 9J3T, resolution: 3.80 Å). It can be seen from the figure that although EM3DFold and ModelAngelo recover comparable nucleotide coverage (84% vs. 81%), EM3DFold achieves much higher sequence accuracy (94% vs. 32%). Further examination reveals that ModelAngelo incorrectly assigns a large fraction of nucleotides as cytosines, while EM3DFold successfully recovers the diverse nucleotide composition. These examples highlight the ability of EM3DFold to build more complete nucleic acid models with improved nucleotide-type assignment in challenging cryo-EM maps.

### 2.3 Performance on 124 protein-nucleic acid complexes

We next evaluated EM3DFold on another independent test set comprising 124 protein-nucleic acid complexes. Figure 3 shows the evaluation results of EM3DFold in comparison with ModelAngelo. It can be seen from the figure that EM3DFold achieves a better backbone RMSD than ModelAngelo (0.75 Å vs. 0.77 Å) (Figure 3a). In particular, EM3DFold significantly improves coverage (95% vs. 84%) (Figure 3b), while also achieving slightly higher sequence accuracy (96% vs. 95%) (Figure 3c). Consequently, EM3DFold achieves a significantly better completeness than MoldelAngelo (90% vs. 79%). It is also noted that for coverage and completeness, EM3DFold consistently outperforms ModelAngelo for nearly all targets (Figure 3d), highlighting its superior ability to construct more complete models.

**Figure 3:**
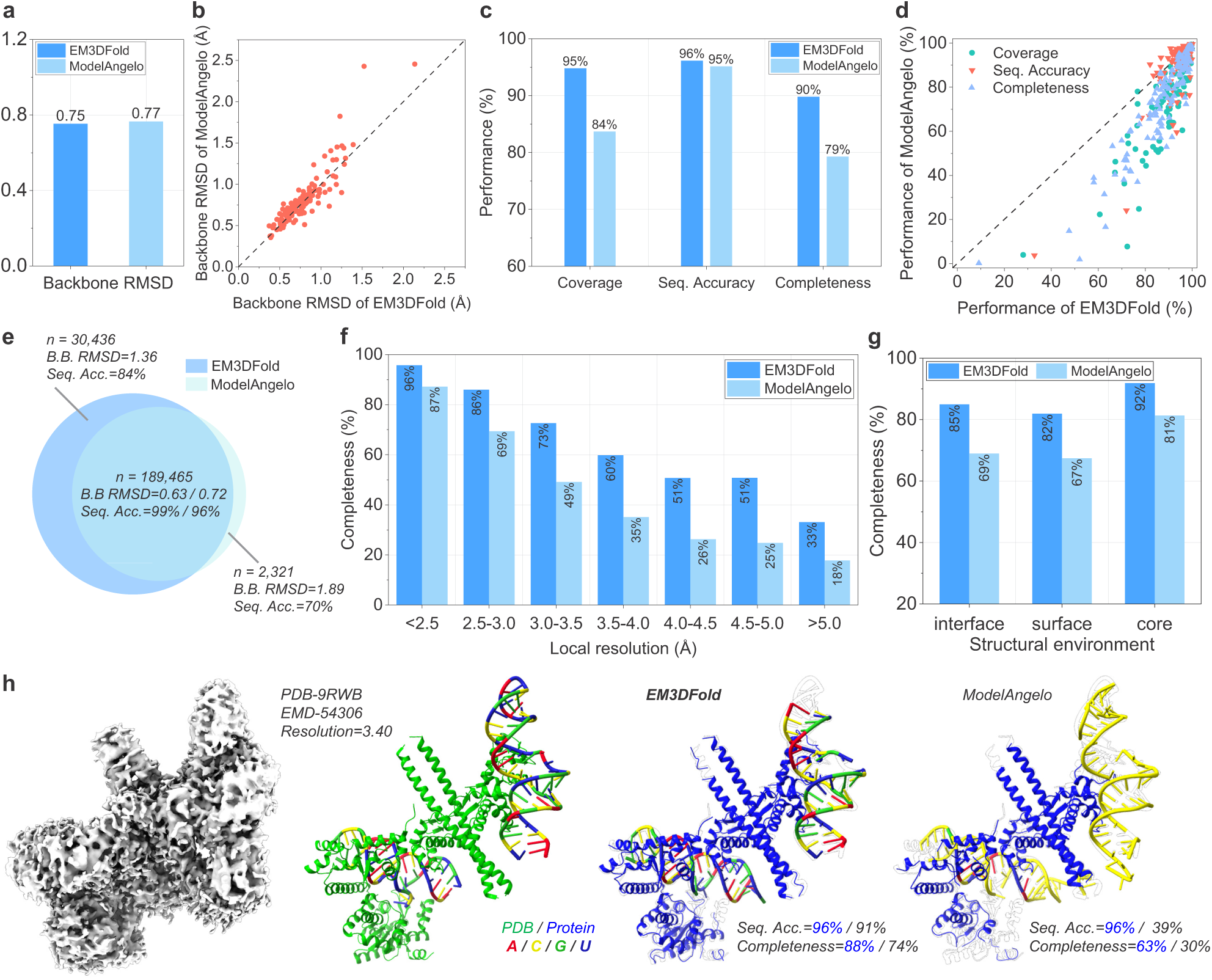
Performance of EM3DFold and ModelAngelo on *n* = 124 protein-nucleic acid complexes. **a**, Backbone RMSDs of EM3DFold and ModelAngelo on protein-nucleic acid complexes (*n* = 124). Bars indicate medians. **b**, Head-to-head comparison of backbone RMSDs between EM3DFold and ModelAngelo. **c**, Bar-plots of coverage, sequence accuracy and completeness for EM3DFold and ModelAngelo. Bars indicate medians. **d**, Head-to-head comparison of coverage, sequence accuracy, and completeness between EM3DFold and ModelAngelo. **e**, Completeness comparison between EM3DFold and ModelAngelo for different local resolution ranges. **f**, Completeness comparison between EM3DFold and ModelAngelo for different structural environments, including interface, surface, and core regions. **g**, Sequence accuracy comparison between EM3DFold and ModelAngelo for different structural environments. **h**, Representative example demonstrating improved protein and nucleic acid modeling by EM3DFold. The PDB structure is shown in green, the EM3DFold model in blue, and the ModelAngelo model in yellow. Nucleotides are colored according to their types. On this target, EM3DFold achieves higher sequence accuracy and completeness for both protein and nucleic acids compared with ModelAngelo.

Figure 3e displays the accuracies of EM3DFold-unique, ModelAngelo-unique, and commonly built residues. Compared with ModelAngelo that uniquely models 2,321 residues with lower accuracy (1.89 Å RMSD and 70% sequence accuracy), EM3DFold uniquely models many more residues (30,436) with a significantly better backbone RMSD of 1.36 Å and sequence accuracy of 84%, demonstrating the high ability of EM3DFold to recover additional regions with reliable structural and sequence assignments (Figure 3e). For the 189,465 commonly built residues, EM3DFold also achieves a better performance than ModelAngelo, with backbone RMSD of 0.63 Å vs. 0.72 Å and sequence accuracy of 99% vs. 96%, respectively (Figure 3e). These results demonstrate that EM3DFold is able to maintain high modeling accuracy while building substantially more residues from challenging regions that are frequently missed by ModelAngelo.

To investigate the robustness of EM3DFold for heterogeneous cryo-EM densities, we evaluated the model completeness in different local resolution ranges (Fig. 3f). It is shown that EM3DFold consistently achieves higher completeness than ModelAngelo across all resolutions, with the advantage becoming increasingly pronounced in lower-quality density regions. For example, at resolutions better than 2.5 Å, EM3DFold achieves a completeness improvement by 11% (98% vs. 87%); however, as the local resolution decreases, the improvement between the two methods progressively continues, e.g. yielding an improvement by 25% completeness for local resolutions of 3.5-5.0 Å. In challenging regions with local resolution worse than 5.0 Å, EM3DFold maintains 33% completeness, whereas ModelAngelo achieves only 18%.

Furthermore, we examined whether EM3DFold maintains its advantage across different structural environments (interface, surface, and core regions) of complex structures (Fig. 3g). It is revealed that EM3DFold also provides more complete models in all structural contexts. Both EM3DFold and ModelAngelo performs better on core regions than other regions as core regions are generally more stable. In core regions, EM3DFold maintains a moderate completeness improvement over ModelAngelo by 11% (92% vs. 81%). The improvement is more evident for exposed regions with an increase by 16% (85% vs. 69%) for interfaces and 15% (82% vs. 67%) for surfaces. These results suggest that EM3DFold is particularly robust in challenging structural environments like interface regions that involve complicated intermolecular interactions.

Figure 3h shows an example of protein-nucleic acid complex (PDB-9RWB, EMD-54306), where EM3DFold achieves a higher completeness for both the protein and nucleic acid components compared with ModelAngelo. On this target, EM3DFold gives a high completeness of 88% and 74% for the protein and nucleic acid components, respectively, compared with 63% and 30% for ModelAngelo.

### 2.4 Validation by model-to-map fit

We further evaluated the map-to-model consistency of built models. Figure 4a-b shows the CC-mask and Q-score values of EM3DFold and other methods on 178 nucleic acid maps. It is shown that EM3DFold outperforms EM2NA, CryoREAD, and ModelAngelo in both metrics. Specifically, for nucleic acid, EM3DFold achieves the Q-score and CC-mask values of 0.55 and 0.61, respectively, compared with 0.49 and 0.51 for EM2NA, 0.18 and 0.18 for CryoREAD, and 0.50 and 0.52 for ModelAngelo (Fig. 4a-b). The head-to-head analysis of Q-scores shows that EM3DFold achieves improved map-model agreement in the majority of test cases (Fig. 4c). Similar trends are also observed in the refined models, although all methods obtain some improvement (Fig. 4a-b.

**Figure 4:**
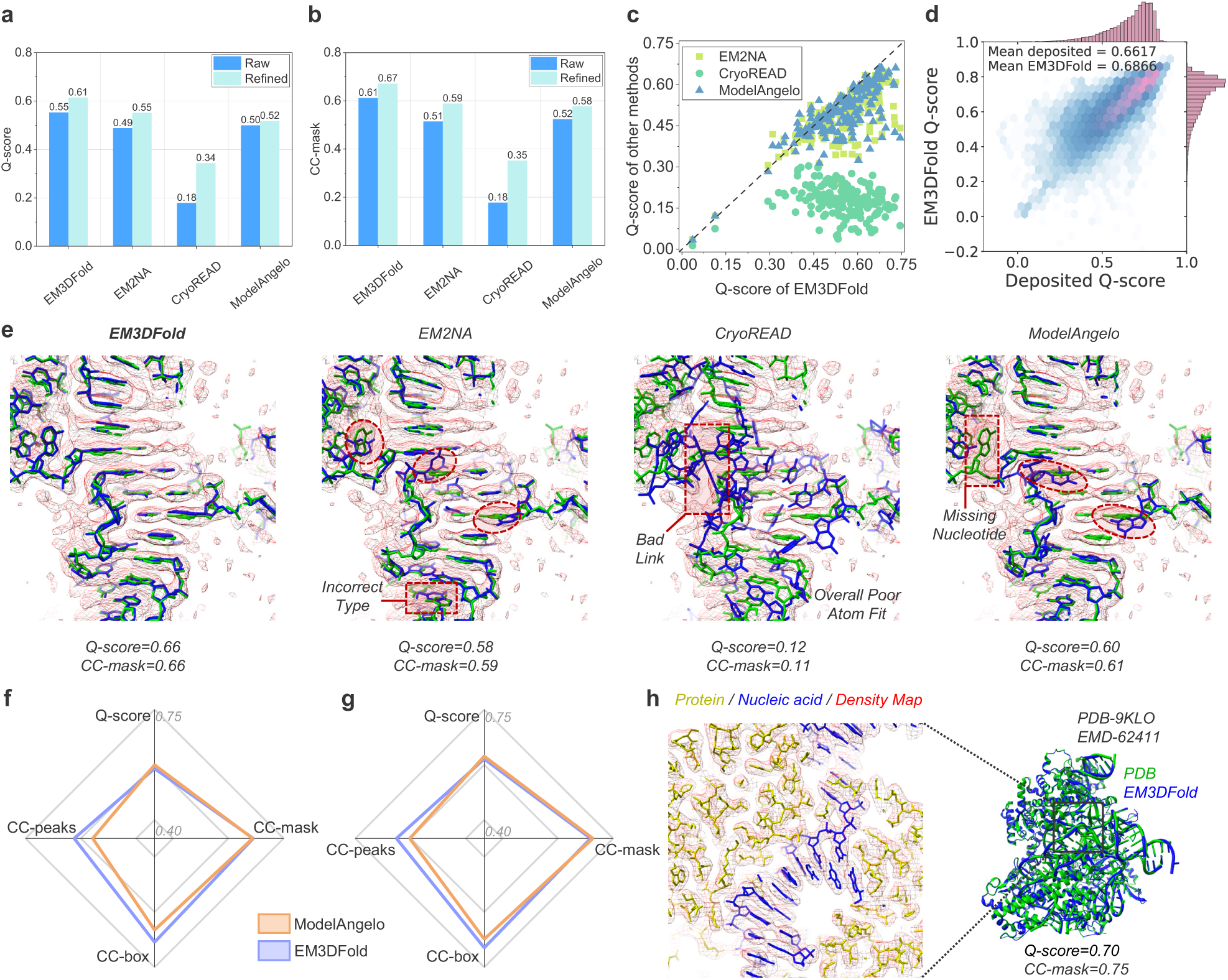
Comparison of EM3DFold and other methods in terms of model-map consistency. **a**, Bar plots of Q-score of nucleic acid models before (Raw) and after Phenix real-space refinement (Refined) (*n* = 178). **b**, Bar plots of CC-mask of nucleic acid models before (Raw) and after Phenix real-space refinement (Refined) (*n* = 178). **c**, Head-to-head comparison of nucleic acid Q-scores between EM3DFold and other automated modeling methods. **d**, Distribution of Q-scores for correctly assigned nucleotides modeled by EM3DFold compared with the corresponding deposited PDB nucleotides. **e**, Representative comparison of nucleic acid modeling by EM3DFold, EM2NA, CryoREAD, and ModelAngelo on the same target (EMD-63014, PDB-9LE3). On this case, EM3DFold accurately builds both the RNA backbone and nucleotide identities. By comparison, EM2NA and ModelAngelo occasionally exhibit modeling errors and atom outliers, and CryoREAD suffers from tracing errors and overall poor density fitting. **f**, Radar plot of average Q-score, CC-mask, CC-box, and CC-peaks, calculated for models before refinement. **g**, Radar plot of average Q-score, CC-mask, CC-box, and CC-peaks, calculated for models after refinement. **h**, Representative protein-nucleic acid complex (EMD-62411, PDB-9KLO) built by EM3DFold. The EM3DFold-built model is colored in green, the reference PDB structure in blue, and the cryo-EM density map in red mesh. Enlarged views illustrate excellent agreement between the predicted model and the experimental map density.

Figure 4d shows the Q-scores of refined EM3DFold models versus those of deposited PDB structures at the residue level. It can be seen from the figure that the Q-scores of the nucleotides built by EM3DFold are strongly correlated with those of the deposited structures. In particular, the EM3DFold models even obtain a higher mean Q-score than the deposited PDB structures (0.6866 vs. 0.6617) (Fig. 4d), suggesting that EM3DFold is able to build more accurate models than expert-level manual model building in terms of atom resolvability^43^.

Figure 4e presents a representative comparison of different methods on the same nucleic acid region. It can be seen from the figure that EM3DFold accurately reconstructs the atomic positions and nucleotide conformations, yielding excellent agreement with both the deposited structure and the experimental density (Q-score=0.66 and CC-mask=0.66). In comparison, although EM2NA and ModelAngelo perform well in most residues (0.58 and 0.59 for EM2NA, 0.60 and 0.61 for ModelAngelo), EM2NA and ModelAngelo both fail on some residues, as highlighted by the dashed circles. CryoREAD exhibits more severe errors, including incorrect backbone connectivity and poor atomic fitting to the density, resulting in substantially lower Q-score (0.12) and CC-mask (0.11).

For protein-nucleic acid complexes, EM3DFold also achieves an overall better performance than ModelAngelo in model-to-map fit (Fig. 4f-g). Specifically, EM3DFold is comparable to ModelAngelo in terms of Q-score for raw (0.59 vs. 0.60) and refined models (0.62 vs. 0.62). However, EM3DFold achieves higher CC-box and CC-peaks values than ModelAngelo for both raw and refined models (Fig. 4f-g). Figure 4h shows a representative protein-nucleic acid complex (EMD-62411, PDB-9KLO) built by EM3DFold. The built model is accurately reconstructed within the cryo-EM density map, achieving an excellent residue coverage/sequence accuracy of 99%/99% for the protein and 92%/95% for the nucleic acid. In addition, individual atoms also fit the map density very well, resulting in a high Q-score (0.70) and CC-mask (0.75) (Fig. 4h). These results demonstrate that EM3DFold achieves superior model-map consistency than state-of-the-art methods for both nucleic acids and protein-nucleic acid complexes

### 2.5 Validation by stereochemical quality

We next assessed the stereochemical quality of built models. This evaluation verifies whether a model conforms to fundamental principles of chemical geometry, including steric clashes, bond lengths, bond angles, torsion angles, etc., without respect to the map. Four metrics are used, including Clash score, MolProbity score, RMS(bonds), and RMS(angles) calculated by the MolProbity program^45^ to quantify geometric rationality. Together with the map-to-model fit analysis, these structural assessments provide a comprehensive measure of model reliability and its readiness for deposition into the PDB.

Table 1 lists the average structure quality results of EM3DFold, EM2NA, CryoREAD, ModelAngelo, and EMProt on 178 nucleic acid and 124 protein targets. It can be seen from the table that EM3DFold obtains substantially improved stereochemical quality for built raw models of nucleic acids. Although all methods benefit from post-refinement, EM3DFold maintains better performance compared with other approaches. This indicates that the improved stereochemical quality of EM3DFold is not solely due to post refinement, but is already present in the initial model generation stage. For protein targets, EM3DFold also shows comparable or better performance in most metrics. Specifically, EM3DFold achieves the best Clash score (Raw), MolProbity scores (raw models), RMS (Bonds) (raw and refined models), and comparable RMS (angles) values (Table 1).

**Table 1:** Stereochemical quality evaluations for the built models by EM3DFold, EM2NA, CryoREAD, ModelAngelo, and EMProt on the main test set containing *n* = 178 nucleic acids and *n* = 124 proteins. Both the results for raw (refined) models are listed.

| Type | Method | Clash score ↓ | MolProbity score ↓ | RMS (bonds) ↓ | RMS (angles) ↓ |
| --- | --- | --- | --- | --- | --- |
| Nucleic acid | EM3DFold | **70.16(18.27)** | **3.43(2.78)** | **0.0575(0.0124)** | **7.57(1.39)** |
|  | EM2NA | 267.52(46.77) | 4.01(3.20) | 0.2475(0.0331) | 13.97(2.04) |
|  | CryoREAD | 248.42(48.60) | 3.99(3.20) | 0.1597(0.0236) | 12.65(4.61) |
|  | ModelAngelo | 217.90(25.83) | 3.93(2.92) | 0.1146(0.0168) | 9.51(1.51) |
| Protein | EM3DFold | **59.99(11.84)** | **3.27(1.98)** | **0.0313(0.0074)** | 4.14(1.06) |
|  | EMProt | 60.36(11.92) | 3.41(1.99) | 0.0314(0.0078) | **4.04(1.01)** |
|  | ModelAngelo | 60.97(**11.13**) | 3.40(**1.92**) | 0.0623(0.0128) | 5.22(1.19) |

The numbers in **bold** indicate the best performances for the corresponding metrics. Downward arrows indicate that lower values are better. Values are reported for Raw (Refined) models.

The evaluation on nucleic acid structure quality highlights a common bottleneck in current modeling approaches, namely the difficulty of simultaneously achieving high structural completeness and stereochemical validity. EM3DFold largely alleviates this issue, enabling more physically realistic all-atom models. Overall, EM3DFold achieves a better balance between stereochemical quality and model completeness, with particularly strong improvements for nucleic acid structures while maintaining competitive performance for proteins.

### 2.6 Further evaluation on 120 protein-only targets

Having demonstrated the capability of EM3DFold in modeling diverse nucleic acid-containing systems, we further assessed its performance on protein-only targets. This evaluation examines whether EM3DFold can provide a general solution for protein structure modeling, as protein complexes still represent the majority of structures determined by cryo-EM. To this end, EM3DFold is evaluated on *n* = 120 protein-only targets and compared with EMProt and ModelAngelo.

Figure 5a-b shows the overall completeness of the built protein models by different methods. As shown in Fig. 5a, EM3DFold achieves the best completeness of 95%, compared with 88% for EMProt and 87% for ModelAngelo. The head-to-head comparison demonstrates that the completeness of EM3DFold is higher than that of the other two methods for nearly all test cases (Fig. 5b). The relationship between model completeness and local resolution is also examined. It is shown that EM3DFold maintains the highest completeness across all resolution ranges (Fig. 5c). The advantage becomes more pronounced at intermediate resolutions. For example, at local resolution of *>* 4.0 Å, EM3DFold still yields a markedly higher completeness of 67%, compared with 24% for EMProt and 30% for ModelAngelo.

**Figure 5:**
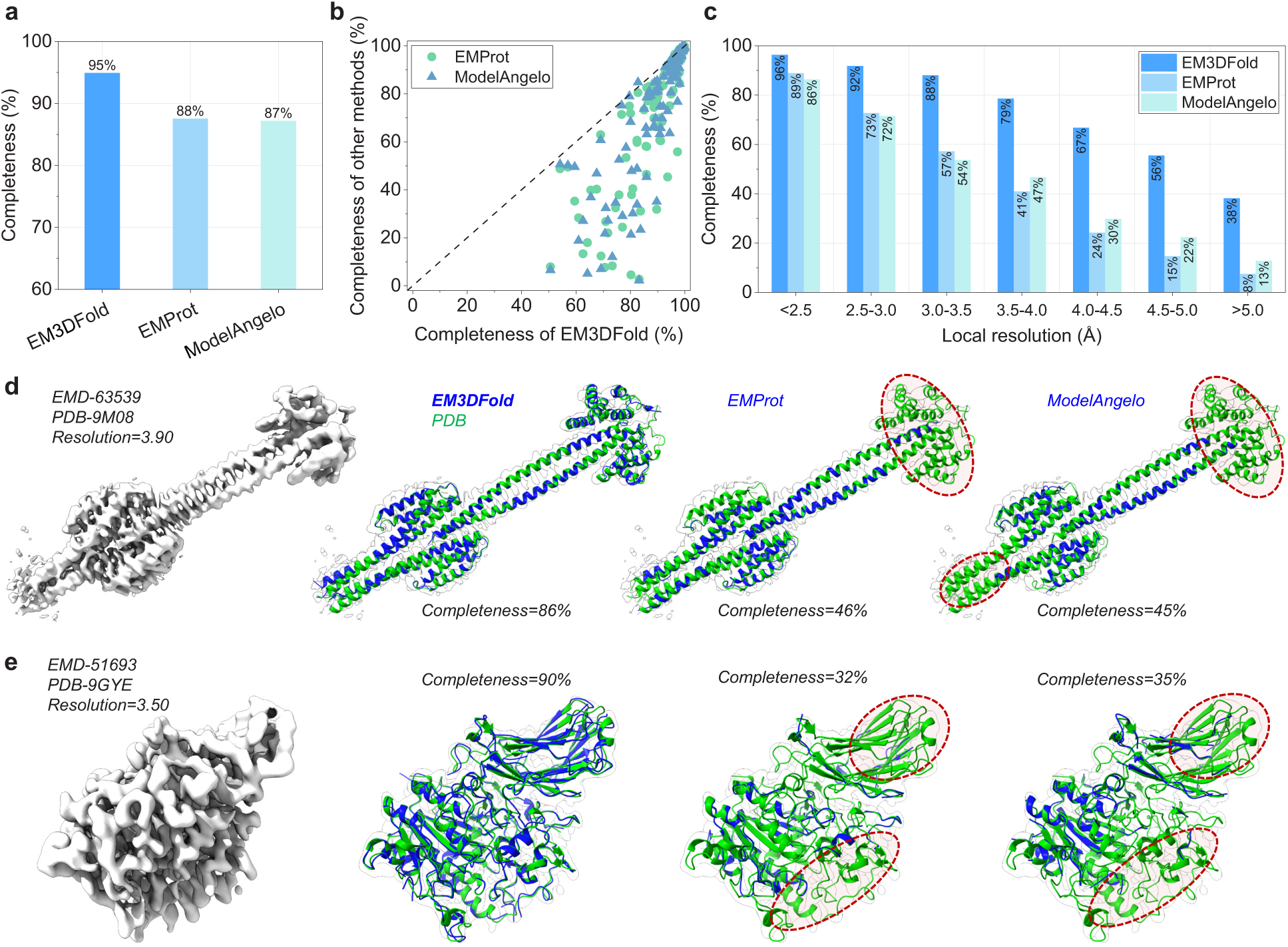
Evaluation of EM3DFold and other methods on *n* = 120 protein-only targets. **a**, Comparison of model completeness between EM3DFold, EMProt, and ModelAngelo on protein-only cryo-EM maps. **b**, Head-to-head comparison of model completeness between EM3DFold and other modeling methods (EMProt and ModelAngelo). **c**, Completeness comparison of EM3DFold, EMProt, and ModelAngelo across different local resolution ranges. **d-e**, Representative examples showing improved model completeness by EM3DFold compared with EMProt and ModelAngelo. The EM3DFold models are shown in blue, and the deposited PDB structures in green. Red dashed circles highlight regions missed by other methods but recovered by EM3DFold. **d**, The structure of outer membrane lipoprotein QseG and histidine kinase QseE complex (PDB-9M08, EMD-63539). **e**, The structure of Hemolytic Phospholipase C (PDB-9GYE, EMD-51693).

Several representative examples are shown in Fig. 5d-e. For target EMD-63539/PDB-9M08 at a resolution of 3.90 Å (Fig. 5d), EM3DFold reconstructs substantially more of the compact domain of the Quorum-sensing regulator protein G, achieving an overall completeness of 86%, whereas EMProt and ModelAngelo only give 46% and 45%, respectively. The missing regions in EMProt and ModelAngelo are mainly located at the N-terminal domain (Fig. 5d). A similar trend is observed for EMD-51693/PDB-9GYE at a resolution of 3.50 Å (Fig. 5e), where EM3DFold achieves 90% completeness, compared with 32% for EMProt and 35% for ModelAngelo. In both examples, EM3DFold produces models that match the deposited structures much more extensively, especially in lower-resolution regions like terminal domain.

### 2.7 Examples of highly accurate models built by EM3DFold

Figure 6 further exhibits the accurately built models by EM3DFold on several representative cases with diverse molecular compositions and architectures. All of these models achieve a high sequence accuracy of *>*85%, indicating that only minor manual adjustment is required in real applications. Here, the demonstrated cases include both small DNA targets of *<*100 nt and very large RNA targets of thousands of nucleotides. These cases also includes protein-nucleic acid, RNA-only and protein-only targets.

**Figure 6:**
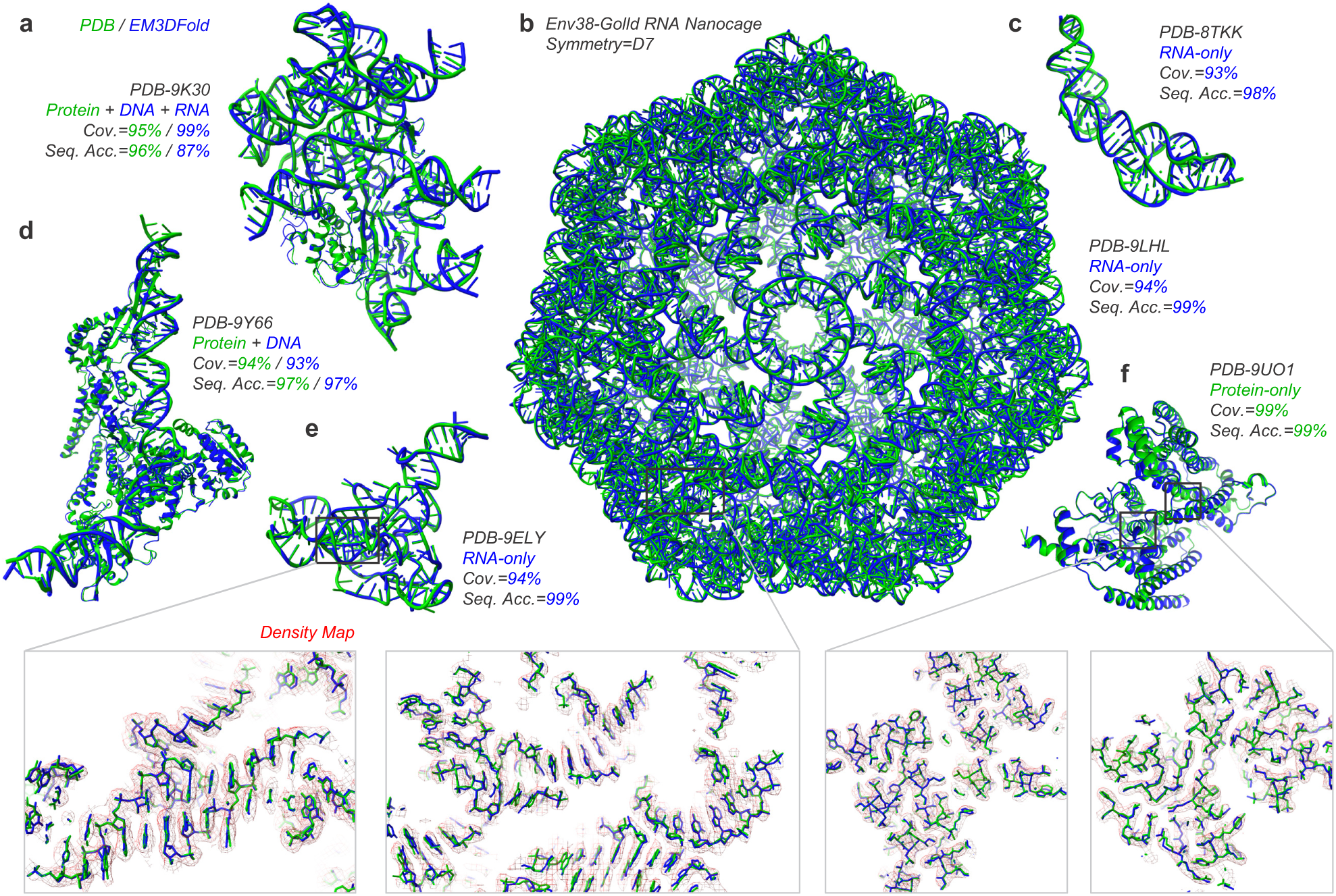
Examples of highly accurate atomic models built by EM3DFold. The EM3DFold models are shown in blue, and the corresponding PDB structures are in green. **a**, The TbaIscB-omega RNA-target DNA complex modeled by EM3DFold (protein-DNA-RNA complex, PDB-9K30). **b**, The Env-38 Golld RNA nanocage exhibiting global D7 symmetry, representing one of the largest RNA multimers determined to date (RNA-only, PDB-9LHL). **c**, The apo-state structure of RNA device 43 truncation mutant 3 (U100C) (RNA-only, PDB-8TKK). **d**, The dimeric AttLsym-bound serine integrase complex (protein-DNA complex, PDB-9Y66). **e**, The RaiA RNA motif (RNA-only, PDB-9ELY). **f**, The human organic solute transporter Ost-alpha/beta bound to DHEAS (protein-only, PDB-9UO1). The bottom panels show enlarged views of the atomic model fitting into the cryo-EM density, with the PDB residues colored green, the EM3DFold-modeled residues colored blue, and the density map shown as red mesh.

As shown in Fig. 6a EM3DFold accurately reconstructs protein, DNA, and RNA mixed ternary complexes (PDB-9K30), achieving coverage/sequence accuracy of 95%/99% and 96%/87% for protein and nucleic acid components, respectively. Figure 6d presents a representative protein-DNA complex (PDB-9Y66), where EM3DFold produces highly complete and accurate models with 97% sequence accuracy for both protein and DNA. Figure 6b shows an example of the RNA nanocage containing up to *>* 10,000 nucleotides^46–49^. The nanocage adopts a global D7-symmetric architecture composed of 14 repeated RNA chains, making complete atomic model building highly challenging. Nevertheless, EM3DFold automatically reconstructed the entire assembly with a high residue coverage of 94%, and a sequence accuracy of 99%, demonstrating accuracy comparable to the deposited PDB structure.

The superior modeling performance of EM3DFold is also illustrated by representative examples shown in Fig. 6c,e. On these targets, EM3DFold accurately reconstructs RNA backbones and assigns nucleotide identities, achieving a residue coverage/sequence accuracy of 93%/98% for PDB-8TKK and 94%/99% for PDB-9ELY. Besides nucleic acid modeling, a protein-only example with 99% completeness is also presented in Fig. 6f. On these targets, the modeled residues show excellent agreement with the cryo-EM density maps, as demonstrated by the high-quality atom-level details enlarged in Fig. 6b,e,f. These representative examples demonstrate that EM3DFold generalizes effectively across diverse macromolecular systems, ranging from compact RNA folds, protein-nucleic acid complexes to large RNA assemblies.

### 2.8 Computational efficiency of EM3DFold

The computational efficiency of EM3DFold was evaluated on several representative RNA-only, protein-nucleic acid, and protein-only targets. All experiments were performed using a single NVIDIA A100 GPU (40 GB) and one CPU core. It is shown that EM3DFold exhibits good computational efficiency across a broad range of target sizes. For structures of median size, EM3DFold completes model building in approximately 10 minutes. For structures containing up to approximately 5,000 residues, model building can be finished within around 30 minutes. The largest RNA-only structure, comprising more than 11,000 nucleotides, was built within 54 minutes. Overall, the runtime increases approximately linearly with the size of the target, demonstrating that EM3DFold maintains practical computational efficiency for routine automated structure determination.

## 3 Discussion

We have proposed a universal density-aware and language model-powered deep learning method for accurate model building of proteins, nucleic acids, and protein-nucleic acid complexes from cryo-EM maps, named EM3DFold. EM3DFold builds the all-atom structure through several steps: C*α*/C4’ atom prediction, all-atom prediction, backbone tracing, residue type prediction, and sequence assignment. The density-aware and language model-powered Three-Track Attention network enables the accurate prediction of residue types and atomic positions by efficiently integrating sequence, density, and structure. The model building process is fully automated and also computationally efficient.

The proposed method was extensively evaluated on independent benchmarks of 298 experimental cryo-EM maps and compared with state-of-the-art automated model-building methods. It is revealed that EM3DFold consistently outperforms existing approaches on nucleic acid, protein, and protein-nucleic acid complex modeling tasks. For nucleic acids, EM3DFold achieves the highest completeness of 75%, substantially outperforming EM2NA (45%), CryoREAD (26%), and ModelAngelo (30%). For protein-nucleic acid complexes, EM3DFold achieves better model completeness (90%) than ModelAngelo (79%). For protein-only targets, EM3DFold also demonstrates significant advantages, achieving the higher completeness (95%) than existed methods (88% for EMProt and 87% for ModelAngelo). Beyond its significant improvements in overall model completeness, EM3DFold also generates atomic models with improved model-to-map fit and stereochemical quality across diverse biomolecular systems.

Despite the present successes, several limitations remain. Similar to other map-based model building methods, the performance of EM3DFold depends on the quality of the input cryo-EM map. Local resolution variations often lead to uneven density quality, which can adversely affect model construction. In particular, weak density regions remain challenging, leading to potential errors in backbone tracing and residue placement. It is also observed that sequence misalignment may still occur in ambiguous regions. In addition, highly symmetric assemblies or regions with repetitive sequence patterns may further complicate sequence assignment. Finally, although EM3DFold generally produces well-formed atomic models, occasional local geometric inconsistencies or overfitting to noisy density regions may still occur. Therefore, the generated models may further benefit from post-refinement using established tools like Phenix or Coot, together with targeted manual intervention, to address local modeling errors and further improve stereochemistry and model-map agreement.

Several directions could be explored to further enhance the modeling capabilities of EM3DFold. First, the incorporation of additional biological priors may improve the robustness in modeling low quality maps. Second, continued scaling of training datasets is expected to further advance the accuracy and generality of nucleic acid modeling. Third, integrating complementary sources of sequence co-evolution signals derived from large-scale multiple sequence alignments (MSA) and modern structure prediction models, may further enhance model accuracy and reliability.

## 4 Methods

### 4.1 Data collection

We searched the PDB and EMDB (as of May 20, 2026) to obtain a complete list of maps and structures with resolution *<* 4 Å. The dataset was further filtered using the following metrics: (1) CC-mask for the protein/nucleic acid structure *>* 0.60; (2) CC-box for the entire structure *>* 0.50; (3) Chimera^50^ correlation between map and model-simulated-map *>* 0.50. We also removed items with other irregularities, e.g., having C*α* atoms only, having many unknown residues etc.

Subsequently, the structures released before January 1, 2025 were kept as candidates for the training set. To maintain a manageable training set, we clustered all sequences using MMseqs2^51^. Structures sharing either a nucleic acid sequence identity greater than 80% or a protein sequence identity greater than 30% were grouped into the same cluster. We then randomly selected up to 10 structures from each cluster. When a cluster contained both nucleic acid-containing targets and protein-only structures, nucleic acid-containing items were preferentially retained. This yields a final training set of 6,870 structures, including 2,528 nucleic acid-containing structures and 4,342 protein-only structures. For training the C*α*/C4’ prediction networks, only up to 1 representative structure was selected from each cluster for training efficiency, resulting in a training set of 1,376 maps. For training the all-atom prediction network, the complete dataset of 6,870 structures were used.

Next, the structures released on or after January 1, 2025 were used as candidates for the test set. To remove the redundancy with the training set, we removed the structures that share either a nucleic acid sequence identity greater than 80% or a protein sequence identity greater than 10% with any target in the training set using MMseqs2. For saving computational time, we further excluded excessively large structures and retained only those containing fewer than 20,000 amino acids or nucleotides. Despite these filtering steps, around 1,000 candidate structures still remains. To obtain an appropriate number of diverse test cases, we then clustered the remaining structures based on sequence similarity (100% for nucleic acids and 30% for proteins) and removed those structures with unknown residues. For nucleic acid-containing clusters, the representatives are selected, forming a total of 178 test samples (main test set). For protein-only clusters, we randomly selected 120 items as test cases. The resulting final data set contains 298 items, consisting of 54 RNA-only structures, 124 protein-nucleic acid complexes, and 120 protein-only structures. The average max-TM-score at the complex level to the training set is 0.24 (calculated by Foldseek^52^), and only 3 RNA families are identical to the training items.

### 4.2 Main-chain atom prediction

#### 4.2.1 The Swin-Conv-UNet network architecture

We adopted the Swin-Conv-UNet (SCUNet)^33^ architecture to extract atom probabilities and nucleotide types from density maps. The choice of such a network architecture is based our previous finding^53^ that SCUNet is superior to conventional UNet architectures in cryo-EM density interpretation tasks. The network comprises three encoder blocks, one transition block, and three decoder blocks, which are all built upon Swin-Conv (SC)^33, 54^ modules with skip connections linking the encoder and decoder stages. Each Swin-Conv block integrates a Swin transformer (SwinT) block in parallel with a residual convolutional (RConv) block, enclosed between two 1 *×* 1 convolution layers. The Swin transformer employs a window size of 3. Down-sampling is performed using a 3D convolution layer with kernel size and stride of 2, while up-sampling is achieved via a 3D transposed convolution layer with the same parameters. The input to the network consists of density chunks of size 48 *×* 48 *×* 48 with a grid spacing of 1.0 Å, and the output has identical dimensions.

#### 4.2.2 Network training

Detection of C*α* and C4’ atoms from experimental density maps is based on a unified SCUNet. Building upon our previous study^31^, EM3DFold adopts a single end-to-end network with two cascaded decoder heads. The first decoder predicts the protein/nucleic acid/background segmentation map from the input density map, while the second decoder predicts the atom probability maps by jointly leveraging the segmentation probabilities and shared encoder features. Specifically,

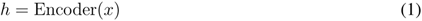

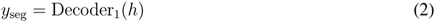

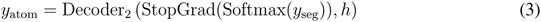

where *x* denotes the input boxes, *h* denotes hidden features. In this way, the predicted semantic segmentation provides explicit structural guidance for atom localization, while both decoder heads are optimized jointly within a single network. We found that directly predicting real-value C*α*/C4’ atom probability using a single decoder led to inferior performance. Without explicit semantic supervision, the network struggles to reliably distinguish protein and nucleic acid atoms, resulting in ambiguous atom localization and reduced prediction accuracy. Introducing the segmentation decoder enables more accurate localization of C*α* and C4’ atoms.

For segmentation supervision, voxels are labeled according to the nearest heavy atom within 3 Å of the reference structure. Each voxel is assigned one of three classes corresponding to protein, nucleic acid, or background. The segmentation loss is defined as a weighted Cross-Entropy loss,

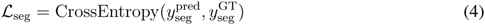

where class weights of 0.25, 0.70, and 0.05 are assigned to protein, nucleic acid, and background voxels, respectively, to alleviate class imbalance.

For C*α* and C4’ prediction, the ground-truth C*α* and C4’ probability maps are generated by

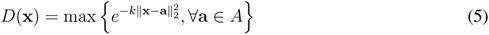

where **a** denotes the coordinate of a C*α* or C4’ atom in the structure *A*. The kernel parameter *k* is determined according to the map resolution *R* as *k* = (*π/*(1.2 + 0.8*R*))^2^, which was originally proposed by DiMaio et al. to maximize the agreement between Gaussian approximations and all-atom density distributions^55^.

All training maps are resampled to a grid spacing of 1.0 Å using trilinear interpolation. Density values are clipped between 0.0 and the 99.999th percentile of each map and then normalized to [0, 1]. Experimental maps and their corresponding labels are split into overlapping chunks of size 60*×*60*×*60 with a stride of 48. Chunks with maximum density values below 0.0 are discarded for computational efficiency. In addition, the chunks with a labeled non-background (protein and nucleic acid) voxel ratio below 0.005 are removed to alleviate the severe imbalance caused by the overwhelming number of background voxels. Finally, approximately 70,000 training chunks are retained.

The atom prediction head is supervised using a combination of Smooth L1 loss and the structural similarity index measure (SSIM) loss^53, 56, 57^

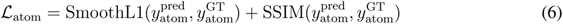

The entire network is optimized end-to-end using the combined multi-task objective,

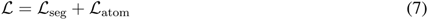

To reduce overfitting, each input chunk is randomly cropped to 48 *×* 48 *×* 48 followed by a random 90*^◦^* rotation. The network is implemented in PyTorch 2.1 and optimized using Adam with an initial learning rate of 5 *×* 10*^−^*^4^ and a weight decay of 1 *×* 10*^−^*^4^. The learning rate is multiplied by 0.95 every two epochs. Training is performed on 2 NVIDIA A100 GPUs (40 GB) with an effective batch size of 48 and typically converges after approximately 200k optimization steps, which takes about 2 days. The model with the lowest validation loss is selected as the final predictor.

### 4.3 All atom construction

According to previous studies, all-atom positions of a residue is reduced into a representative atom, a local backbone frame and torsion angles. Here, for nucleotides, we select C4’ atom as the representative atom, and a local frame is defined with extra C3’ and O4’ atoms. For amino acids, we select C*α* as the representative atom, and N-C*α*-C as the local frame. The positions of the remaining atoms are determined by transforming ideal rigid atom groups according to the torsion angles, thereby constructing the all-atom model.

After obtaining the C*α* and C4’ atom probability map, the C*α* and C4’ atoms are identified as local maxima using a mean-shift algorithm. A Three-Track Attention (TTA) network is then used to predict backbone frames (*R,* **t**) and torsion angles *τ*, given the C*α* and C4’ atoms as inputs. The network comprises 12 main blocks, which iteratively process 1D, 2D, and 3D features.

#### 4.3.1 The Three-Track Attention network architecture

For the 1D input, the densities around each C*α*/C4’ atom (a cubic region of 23 *×* 23 *×* 23 Å^3^) are extracted and then embedded by a convolutional layer. For the 2D input, the pairwise densities between C*α* atoms (a cuboid of 17 *×* 3 *×* 3 Å^3^) are extracted and also embedded by a convolutional layer. Feature updates follow three interaction pathways: (1) 1D features are updated by self-attention and invariant point attention, with biases from 2D features and 3D frames; (2) 2D features are updated from 1D features using outer products; (3) 2D features are updated by axial attention with biases from 3D frames; (4) 3D backbone frames are updated using the updated 1D and 2D features in the structure module. For different prediction tasks, including pRMSDs, torsion angles, and residue types, simple linear layers are employed as prediction heads.

The TTA network here shares a similar concept to the graph network in ModelAngelo^19^, that is, both methods incorporate local cryo-EM density features into the network. However, there are major differences in their overall architecture design. Specifically, EM3DFold explicitly models both node and edge representations, whereas ModelAngelo mainly relies on node representations. In particular, we disentangle cryo-EM density feature extraction into a density initialization module followed by iterative node and edge update modules, rather than integrating these operations within a single CryoAttention module as in ModelAngelo. As a result, our architecture more closely resembles the design of AlphaFold2^40^ and/or RoseTTAFold^58^ by performing attention-based information propagation over both node and edge representations. In addition, we replace ESM-1b with the more recent ESM-2^36^, which has been shown to provide improved representations for structure-related tasks. For nucleic acids, we further incorporate sequence embeddings from the recently developed RiNALMo^37^, enabling the TTA network to exploit contextual sequence information for more accurate nucleotide-type prediction.

#### 4.3.2 Network training

Training follows a similar noising and denoising strategy to that used in ModelAngelo^19^. For each residue, the C*α*/C4’ position **t***^′^*is perturbed by Gaussian noise:

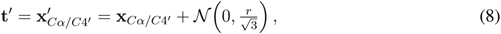

where 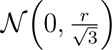 denotes a Gaussian distribution applied to the true coordinate. For nucleic acids, *r* is set to 1.0. For nucleic acid residues, we used a slightly larger value of *r* = 1.2 to account for their higher RMSDs. The rotation matrix *R^′^* of the backbone frame is noised as

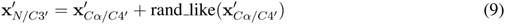

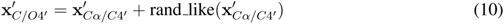

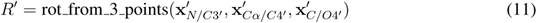

Each training step involves four stages: (1) choosing a residue from structure, (2) extracting its *k* = 200 neighboring residues, (3) applying Gaussian noise, and (4) predicting the true structure with the network. During training, 0-20% of residues are randomly deleted, and 0-20% are replaced with short fragments consisting of 2-5 residues to simulate realistic C*α*/C4’ predictions.

The loss function combines multiple learning objectives. The most important one is the RMSD loss, defined by C*α*/C4’, backbone atoms, and all atoms. The C*α*/C4’ and backbone RMSD losses are computed after every Backbone Update layer, with the *n*-th block weighted by *γ^N−n^* (*γ* = 0.80 in this study) as follows.

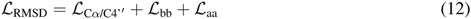

Besides, a torsion angle loss *L*_torsion_ is used to supervise torsion angle predictions, a structure violation loss *L*_violation_ is used to penalize implausible model geometry, a *L*_pRMSD_ loss is used to supervise the pRMSD, and a residue category loss *L*_res-type_ is used for residue typing. We also used some auxiliary losses *L*_aux_ to help the model learn local structure arrangement, e.g. a bond loss *L*_bond_ to supervise whether two nodes have direct C-N/O3’-P bond linking, and a category loss *L*_ss_ to identity protein secondary structures. The total loss is a combination of the above losses as follows.

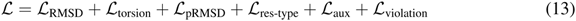

where the violation loss *L*_violation_ is only enabled in the fine-tuning stage.

Training was performed on 4 NVIDIA A100 GPUs (40 GB each) with an effective batch size of 4 for a maximum of 150k steps. Model was trained in two stages: 120k steps for initial training and 30k steps for fine-tuning, which takes about 4 days. The only difference between the initial training and fine-tuning is the inclusion of structure violation loss. Adam optimizer was used for training. The initial learning rate is 2 *×* 10*^−^*^4^ and was multiplied by 0.95 every 2 epochs throughout training. Gradient stopping is applied on rotations between consecutive blocks to stabilize training. Random 1-4 steps are sampled for recycling during training. For inference, the recycling step is set to 4.

### 4.4 Backbone tracing

For each C*α*/C4’ atom, after the local frames and torsion angles are predicted by the all-atom refinement module, backbone atom positions can be determined. Since the nucleotides of the backbone are initially unordered, they must be connected into chains. Residues are iteratively threaded by selecting the next nearest spatial neighbor of the current nucleotide. A chain is terminated and a new one is initiated if two consecutive nucleotides exhibit an O3’–P bond length of *>* 2.5 Å amino acids exhibit a C–N bond length of *>* 2.1 Å and two consecutive

### 4.5 Residue type prediction

Two types of networks, the TTA-based and the SCUNet-based, are trained to predict the residue types in EM3DFold. For proteins, the TTA-based network is used, as described above (Section 4.3.1). For nucleic acids, besides the TTA-based network, we also train a SCUNet-based network following the strategy of EM2NA^31^, as it can yield good nucleotide type prediction results on some high-resolution maps. Combining the Three-Track Attention network with the SCUNet-based predictor achieves better performance for nucleic acids than using either model alone. Specifically, to train this SCUNet type predictor, the PDB model is used to generate labeled type maps: for a voxel, if its nearest heavy atom is within 3 Å it is labeled as the type of the nucleotide (0/1/2/3 for A/C/G/U and 4 for other situations) that the heavy atom belongs to. The same box splitting, normalization, and data augmentation used in C*α*/C4’ atom prediction is also adopted here. The hyper parameters are kept the same as those used for C*α*/C4’ prediction.

### 4.6 Sequence assignment

With the predicted residue types, different sequence assignment strategies are adopted for proteins and nucleic acids. Similar to previous work^19^, for proteins, residue-type prediction from the TTA network is sufficiently accurate and is therefore directly used to construct the HMM profile for each backbone fragment. For nucleic acids, two HMM profiles are constructed, one from the TTA predictions and the other from the SCUNet predictions. Each profile is searched against the target sequence using HMMER^38, 39^, and the alignment with the higher HMM score is retained. This hybrid strategy exploits the complementary strengths of the two residue-type predictors, leading to more robust nucleotide sequence assignment. The selected alignment is then used to assign amino-acid or nucleotide identities to the corresponding fragment. Since it is hard to determine which part of sequences remain unsolved in raw density maps before a model is built, we use the full-length PDB SEQ RES record as target sequences.

### 4.7 Evaluation of built models

Four primary metrics, including backbone RMSD, residue coverage, sequence accuracy, and completeness, are used to evaluate the accuracy of built models. Backbone RMSDs are calculated based on backbone atoms shared among different amino acid or nucleotide types. Specifically, the backbone atoms include N, C*α*, C, and O for proteins, and phosphate atoms (P, OP1, and OP2) together with ribose atoms (C1’, C2’, C3’, C4’, C5’, O3’, O4’, and O5’) for nucleotides. Residue coverage quantifies the fraction of residues in the built model that lie within a distance cutoff from the reference PDB structure, regardless of residue type. Sequence accuracy measures the fraction of residues assigned with the correct residue type. During evaluation of sequence metrics, DA/DC/DG/DT are considered equal to A/C/G/U. Completeness is the product of residue coverage and sequence accuracy, which is to measure the overall performance of a method in model building.

To assess the model-map consistency of built models, another four metrics, including Q-score^43^, CC-mask, CC-box and CC-peaks are used. Here, the Q-score is a measurement of the ‘atom resolvability’ in cryo-EM maps. The CC values measure the correlations between the built model and the map density. All four metrics indicate how an atomic model conforms to the map. It should be noted that these four metrics primarily evaluate model-to-map consistency rather than sequence assignment accuracy, which is assessed independently using sequence accuracy and completeness.

To assess the stereochemical quality of built models, additional four metrics including Clash score, MolProbity score, RMS(bonds), and RMS(angles), which are calculated by phenix.molprobity^45^ program, are also reported. In addition, initially built models are often refined by a third-party program to further improve geometric structure quality and model-map consistency in real applications. Therefore, we have also reported the results of the refined models after post-refinement using the phenix.real space refine tool^59^ in the Phenix package^7^.

## Code availability

The EM3DFold package is freely available at https://github.com/huang-laboratory/EM3DFold.

## Acknowledgements

This work was supported by the National Natural Science Foundation of China (grant Nos. 32430020, 32161133002, and 62072199), the Postdoctoral Fellowship Program of CPSF (grant No. GZB20250617), and the startup grant of Huazhong University of Science and Technology.

## Author contributions

S.H. conceived and supervised the project. T.L. designed and performed the experiments. S.H. and T.L analyzed the data. T.L. and H.C. tested the program. T.L. and S.H. wrote the manuscript. All authors reviewed and approved the final version of the manuscript.

